# Atomic resolution Protein Allostery from the multi-state Structure of a PDZ Domain

**DOI:** 10.1101/2021.11.23.469649

**Authors:** Dzmitry Ashkinadze, Harindranath Kadavath, Celestine N. Chi, Michael Friedmann, Dean Strotz, Pratibha Kumari, Martina Minges, Riccardo Cadalbert, Stefan Königl, Peter Güntert, Beat Vögeli, Roland Riek

## Abstract

Recent methodological advances in solution NMR allow the determination of multi-state protein structures and provide insights into correlated motion at atomic resolution as demonstrated here for the well-studied PDZ2 domain of protein human tyrosine phosphatase 1E for which protein allostery was predicted. Two-state protein structures were calculated for both the free form and in complex with the RA-GEF2 peptide using the exact nuclear Overhauser effect (eNOE) method. In the apo protein states an allosteric conformational preselection step comprising almost 60% of the domain was detected with an “open” ligand welcoming state and a “closed” state that obstructs the binding site by the distance between the β-sheet, α-helix 2 and sidechains of residues Lys38 and Lys72. Observed apo-holo structural rearrangements of induced fit-type are in line with previously published evolution-based analysis covering ~25% of the domain with only a partial overlap with the protein allostery of the open form. These presented structural studies highlight the presence of a dedicated highly optimized dynamic interplay of the complexity of the PDZ2 domain owed by the structure-dynamics landscape.

## INTRODUCTION

An important area of structural biology is the elucidation of the molecular mechanisms behind enzyme activity, protein target recognition and transduction of biological signals enabling various cellular pathways, that rely on protein fold and dynamics (Ishima & Torchia, 2000). Correlated protein motion spanning across the scaffold is of particular interest as it gives rise to protein states and equilibrium (Bai & Englander, 1996; Karplus & Weaver, 1976). One of the most studied types of such conserved protein motion is a ligand-induced protein rearrangement that enables to transduce the signal through a biomolecule in a manner known as allostery. There are two broadly accepted allosteric models including population-shift model and dynamic allostery model (Cooper & Dryden, 1984; Monnot *et al*, 1996). The population-shift model explains protein motion based on the protein population equilibrium between distinct protein conformations whereas dynamic allostery relies on statistical thermodynamics that can explain dynamics-based signal transmission without a change in the average structure.

Typically, allostery is structurally studied indirectly for large systems including enzymes or multimeric proteins using X-ray crystallography of distinct states (Swain & Gierasch, 2006). NMR studies were able to show by chemical shift or relaxation probes allosteric communication within a protein fold in solution lacking however atomic resolution (Emmanouilidis *et al*, 2017; Frauenfelder *et al*, 2001; Fuentes *et al*, 2004; Green & Shortle, 1993; Peng, 2015; Rod *et al*, 2003; Volkman *et al*, 2001). Straight forwardly the ligand binding-induced chemical shift changes (for example measured in a [^15^N,^1^H]-correlation experiment) captures ligand-induced protein scaffold rearrangements and allosteric interactions (Selvaratnam *et al*, 2011). As chemical shifts are more sensitive to the direct interaction with the binding partner, shifts in the protein binding site dominate usually allosterically invoked shifts.

The recent advances in biological NMR allow us to gain insights into the protein motion at atomic resolution by determining multiple protein states using NMR supplied with a plethora of experimental restraints including residual dipolar couplings (RDC), cross-correlated relaxation (CCR), paramagnetic relaxation enhancement (PRE), and Nuclear Overhauser Effect (NOE) restraints (Clore *et al*, 1999; Fenwick *et al*, 2016; Güntert, 2004; Güntert & Buchner, 2015; Güntert *et al*, 1997; Iwahara *et al*, 2004; Kumar, 1985; Vögeli & Vugmeyster, 2019; Wüthrich, 1990). In particular the exact Nuclear Overhauser Effect (eNOE) can yield experimental distances with resolution of up to 0.1 Å (Nichols *et al*, 2017; Orts *et al*, 2012; Strotz *et al*, 2017; Vögeli, 2014; Vögeli *et al*, 2016) that together with the structure calculation of multiple protein states, correction of spin diffusion with eNORA and an automated implementation in the eNORA2 package within CYANA allows a straight forward execution of eNOE based structure calculations (Nichols *et al*, 2018; Orts *et al.*, 2012; Strotz *et al.*, 2017; Vögeli *et al*, 2012). Furthermore, correlated protein motion can be studied using the distance and angle statistics of calculated protein conformers (Ashkinadze *et al*; Fenwick *et al*, 2011a; Fenwick *et al*, 2011b; Fenwick *et al*, 2014).

One of the most studied family of allosteric molecules are PDZ domains (Petit *et al*, 2009). These domains are crucial for protein-protein recognition and protein complex assemblies in multicellular organisms (Tonikian *et al*, 2008). PDZ domains generally recognize the carboxyl-terminus of various target proteins and take part in many cellular processes including cell growth and proliferation. PDZ domains display a compact fold out of six β-strands, two α-helices and a unique flexible loop at the bottom of the binding pocket (Walma *et al*, 2002). The strands of the protein form an antiparallel β-sheet that serves as a platform for target molecule binding. Human tyrosine phosphatase 1E protein (hPTP1E) contains a PDZ2 domain and mediates a series of crucial biological processes such as protein-protein interaction (Harris & Lim, 2001; Hung & Sheng, 2002), signaling (Palmer *et al*, 2002) and apoptosis (Sato *et al*, 1995). The C-terminal peptide derived from the Ras-associated guanine nucleotide exchange factor 2 (RA-GEF2) is a versatile binding partner of the PDZ2 domain of hPTP1E (Harris & Lim, 2001). Solution NMR structures of the PDZ2 domain were solved for a free form as well as for the form bound to the RA-GEF2 (Kozlov *et al*, 2002; Kozlov *et al*, 2000). The PDZ2 domain of hPTP1E binds RA-GEF2 by a β-strand addition between β-strand 2 and α-helix 2 similar to other PDZ domains (Doyle *et al*, 1996). The PDZ2 allostery was studied with various techniques including the use of molecular dynamics (Dhulesia *et al*, 2008; Gerek & Ozkan, 2011), evolutionary data (Lockless & Ranganathan, 1999) and NMR relaxation of protein methyls (Fuentes *et al.*, 2004).

In this study we showcase the use of the recently emerged eNOE approach to investigate correlated motion, induced-fit ligand binding mode, and protein allostery enabled by the structural information of individual protein states at atomic resolution of both free and bound forms of the PDZ2 domain of hPTP1E.

## RESULTS AND DISCUSSION

### Ligand-induced dynamic changes of PDZ2 domain

Heteronuclear 2D NMR spectroscopy was applied in order to gain a qualitative understanding of the PDZ2 domain of hPTP1E binding to the C-terminal peptide derived from the Ras-associated guanine nucleotide exchange factor 2 (RA-GEF2; Ac-ENEQVSAV-COOH) and allosteric interactions. A [^1^H,^15^N]-HSQC spectrum was acquired at 298 K for a uniformly enriched ^15^N-labeled PDZ2 domain both for a free form and bound to the peptide supplied in a two-fold excess. An overlay of the [^1^H,^15^N]-HSQC spectra together with a chemical shift perturbation (CSP) mapped on the protein 3D structure are shown in Figure 1. Averaged atom-weighted chemical shift perturbations were mapped on the later calculated apo protein structure to delineate the allosteric system of the PDZ2 domain. Absolute CSP values are shown in the Figure S1. In line with previous reports residues of the PDZ2 binding site showed significant chemical shift perturbations together with a number of allosteric residues as expected from a highly dynamic scaffold (Fuentes *et al.*, 2004; Walma *et al.*, 2002). In detail, significant CSPs are observed for residues at the β-strand 2 and α-helix 2 which sandwich the ligand upon binding with less prominent CSPs in the flexible loop Gly24-Gly34, β-strand 3, and α-helix 1 of which the latter two are far away from the binding site and thus have been identified as allosteric sites (Fuentes *et al.*, 2004).

**Figure 1:**
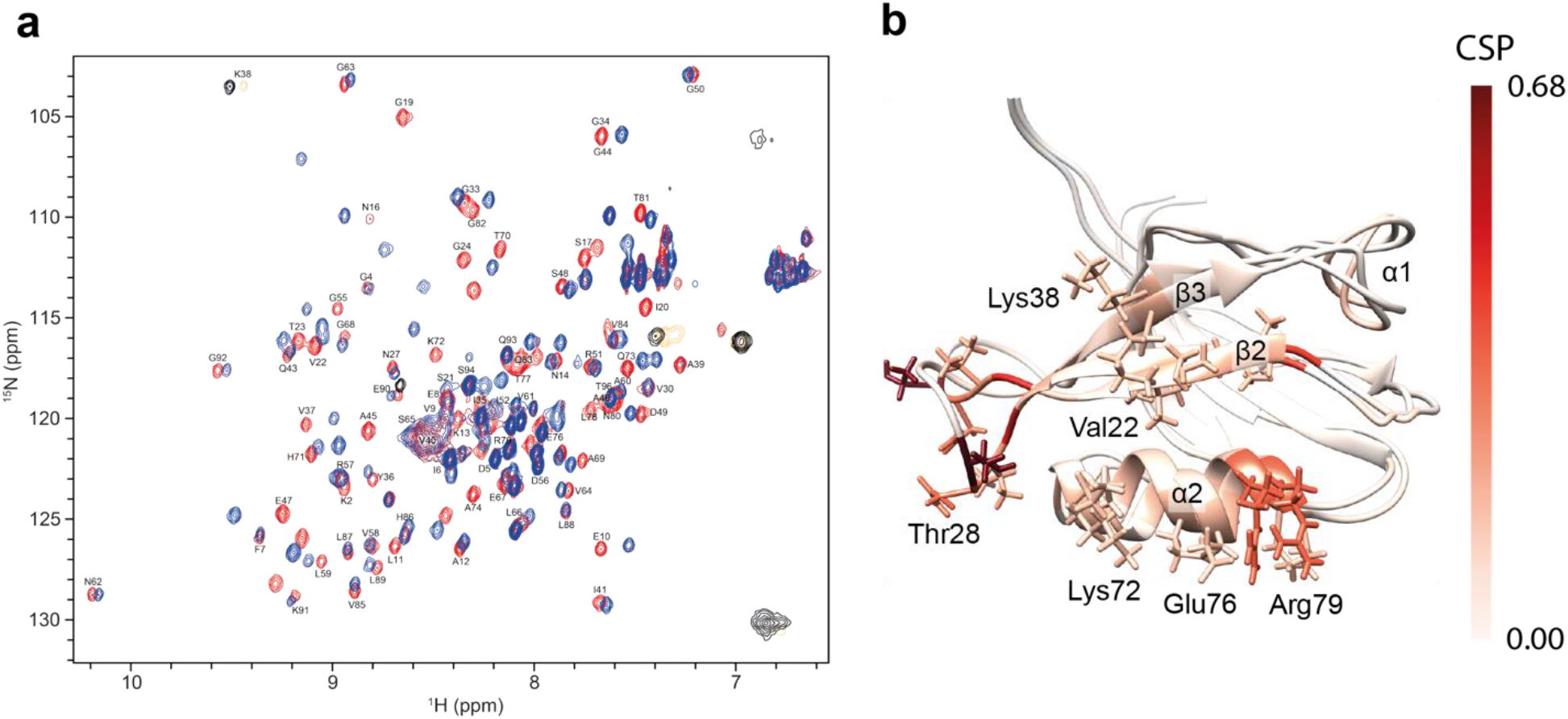
Ligand-binding induced conformational changes measured by chemical shifts. [^1^H,^15^N]-HSQC spectra for the PDZ2 domain in apo form (red) and bound to the RA-GEF2 peptide (blue) (a). Residues of the two-state PDZ2 apo structure with lowest CYANA target function were colored according to the mean of the chemical shift perturbation of ^15^N and ^1^H (CSPs) as indicated by the bar on the right (b).

### Multi-state structure determination of the PDZ2 domain

For apo PDZ2 experimental restraint collection included 1553 distance restraints from eNOEs extracted from a set of 3D [^15^N,^13^C]-resolved [^1^H,^1^H]-NOESY-HSQC spectra at 8, 16, 24, 32, 40, 48 and 80 ms NOESY mixing times (with 410 bidirectional distance restrains with highest precision). In addition, 130 scalar couplings were collected that resulted in ~17 restraints per residue as summarized in Table S1. Similarly for the complex PDZ2 1484 distance restraints from eNOEs and 65 scalar couplings that resulted in ~16 restraints per residue were collected as summarized in Table S2.

In general, with this experimental input multi-state structure calculation with the program CYANA can be performed. CYANA simultaneously optimizes multiple structural states by minimization of the target function calculated by comparing the simulated distance back calculated from all optimized states with experimental upper and lower limit restraints extracted from the NOE cross-peak. Individual states are kept in proximity of each other by symmetry restraints, but local movements with amplitude below 1.2 Å are allowed. Such an approach provides an additional flexibility to the protein fold and allows to observe concerted protein motion. Spin diffusion is a major factor causing inaccuracy in deriving distances from the NOEs (Keepers & James, 1984). Theoretically, spin diffusion can be accounted for if all NOE cross and diagonal peaks can be measured unambiguously, which is unrealistic given a limit to NMR sensitivity (Dalvit *et al*, 2001). Therefore, an alternative version of the spin diffusion correction referred to as exact NOE by Relaxation matrix Analysis (eNORA) was implemented as a part of the eNOE technique that relies on a given 3D protein structure (Orts *et al.*, 2012). So far eNOE analysis has successfully been applied to the proteins WW domain of Pin1, full-length Pin1, GB3, cyclophilin A, ubiquitin and 14-mer UUCG tetraloop (Born *et al*, 2021; Chi *et al*, 2015a; Nichols *et al.*, 2018; Strotz *et al*, 2020; Vögeli *et al.*, 2012; Vögeli *et al*, 2009), and a small RNA tetraloop. Furthermore, a program PDBcor was developed to elucidate correlated motion in form of structural correlations in an unbiased and automated way from the distance statistics of individual structural entities in a multi-state structure (Ashkinadze *et al.*). It can quantify correlations in structural ensembles, uncover the protein regions that undergo synchronized motion and optimally separate conformers into states. PDBcor uses information theory and systematic clustering of protein conformers with the aim to extract mutual information between individual residues (Ashkinadze *et al.*).

In the case of the PDZ2 domain both apo and holo multi-state protein structures were calculated following the established protocol introduced above (Chi *et al.*, 2015a; Strotz *et al.*, 2020; Vögeli *et al.*, 2012) using eNORA2 for the spin diffusion correction and distance extraction (Orts *et al.*, 2012; Strotz *et al.*, 2017) and CYANA for the protein structure calculation with minor modifications (Güntert *et al.*, 1997). Symmetry restraints that keep structural entities in proximity of each other were relaxed in the region starting from the Gly24 up to the Gly34 in order to allow for the additional motion amplitude for the PDZ2 flexible loop. The flexibility in the loop region was shown with R__2__/R__1__ NMR relaxation rates (Figure S2). The structure annealing algorithm was executed with 100’000 energy minimization steps for 1000 two-state conformers. A series of 1-9 state structure calculations were performed (Figure S3) indicating that a single-state structure does not fulfill the experimental data well due to its high CYANA target function (TF), which is a measure of restraint violations, while two states appear to be sufficient to describe the experimental data (Figures S3, S4 and S5). However, the relative population of the two states could not be identified with the use of the CYANA target function due to the low target function contrast for both apo and holo PDZ2 forms (Figure S6).

The twenty two-state conformers with the lowest TF were selected to represent the calculated two-state structure. They satisfy the experimental restraints well as indicated by the low CYANA target function (see Tables S1 and S2) and show favorable Ramachandran plot statistics with less than 2% of the residues in the disallowed regions (Table S3 and Figure S7). In addition, the resulting structures reproduce the known PDZ2 protein fold with root mean square deviations of 1.11 Å for apo and 1.34 Å for the complex from the corresponding reported crystal structures (Tables S1 and S2). Following a jack-knife procedure using the PDBcor software it was determined that correlations presented in the following are experimentally overdetermined in the two-state apo structure as ~70% of the experimental distance restraints are required for the emergence of significant structural correlations (Figure S8).

### The two-state structures of the PDZ2 domain of the apo and holo forms

The eNOE-based two-state structures of the free PDZ2 domain (apo) and in complex with the peptide RA-GEF2 (holo) represented by twenty conformers for each state comprise overall the expected PDZ fold (Figure 2a, 2b, 2d and 2e). First, we analyzed the two states with the PDBcor software in standard settings (Ashkinadze *et al.*) (Figure 2c and 2f). In apo, and in part also in the holo form, we were able to separate the states for the β-sheet, α-helix 2, and the flexible loop Gly24-Gly34. For both apo and holo forms, two states are separated as Chi1 angle values of all calculated protein conformers are split between two state-dependent values reminiscent of local distinct configurations of the side chains (see Figures S3, S4). In both apo and holo PDZ2 forms the flexible loop adopts two conformations, one of which is relatively loose and further apart from the binding site (dark blue state for the apo and red state for the holo form in Figure 2) and another one is more confined and closer to the binding site (cyan state for the apo and yellow state for the holo form; note for convenience the conformer separation and coloring will be kept consistent throughout the manuscript and we will refer to the state with the more confined flexible loop that is closer to the binding site as state 1 and the other state as state 2, respectively).

**Figure 2:**
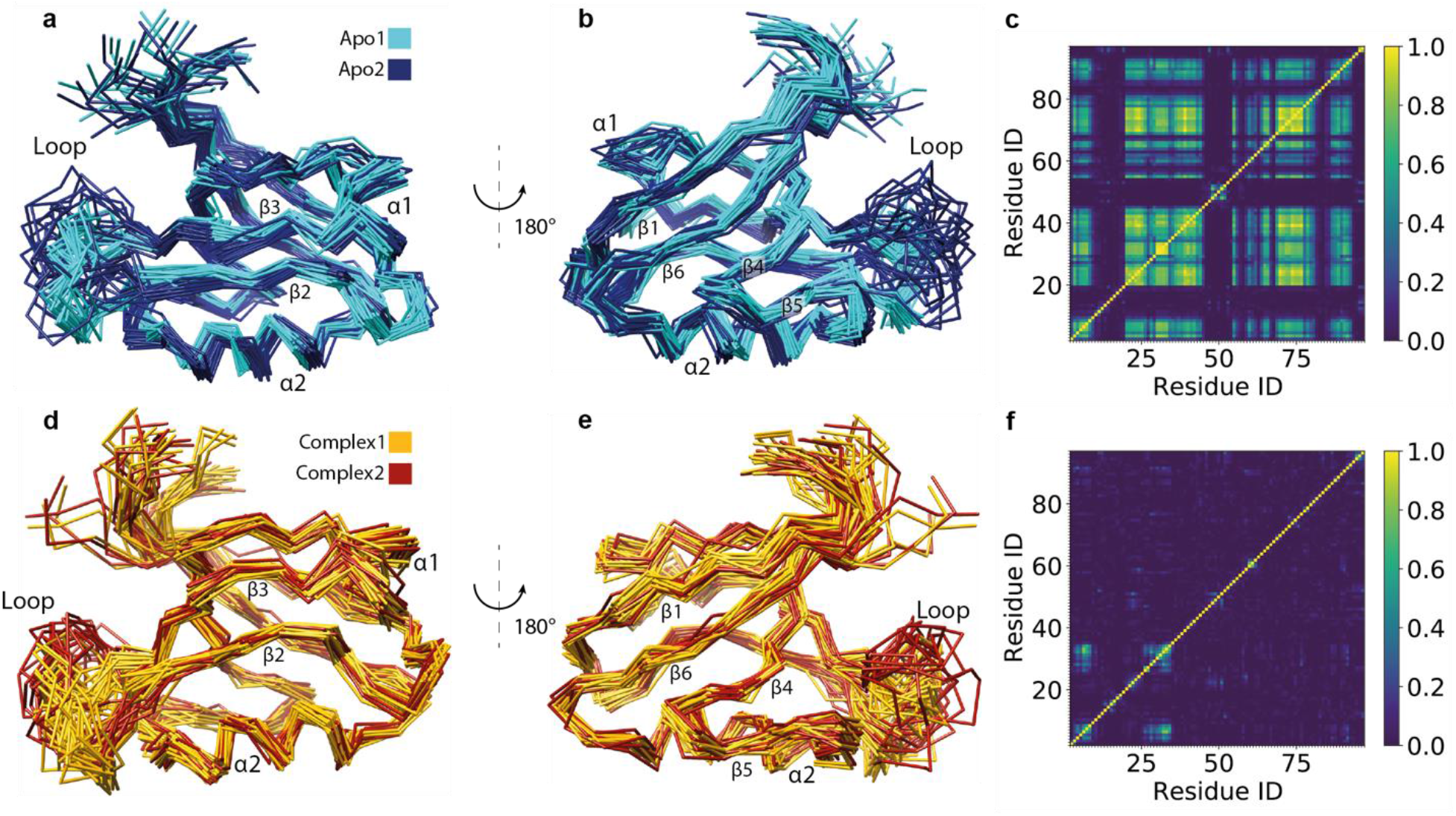
Two-state ensemble structures of the PDZ2 domain in the ligand-free form (a, b) and bound to the RA-GEF2 peptide (d, e) calculated with eNORA2 (Orts *et al.*, 2012; Strotz *et al.*, 2017) and CYANA software in two different orientations. Two apo states are colored in cyan and dark blue whereas two holo states are colored in yellow and red. The secondary structures and the loop comprising residues Gly24-Gly34 are indicated. Structural correlations and optimal state separations for both protein ensembles were calculated with PDBcor in standard settings (Ashkinadze *et al.*) and shown as distance correlation matrix heatmaps for the apo (c) and holo (f) forms of the PDZ2 domain.

The two separate states for the apo form are clearly separable by means of both visual inspection (Figure 2a and 2b) and the correlation map of the PDBcor (Figure 2c). Almost 60% of the entire protein domain appears to shuffle between two states with the most prominent structural difference around the peptide ligand binding site comprising β-strand 2, α-helix 2 and the loop spanning residues Gly24-Gly34 but to a smaller degree extending also into the entire β-sheet, as to be discussed in details below. Structural correlations of the holo PDZ2 domain are much less pronounced. While there are local correlations of the holo states, no global correlation is present (Figures 2f and S4).

### Ligand induced conformational rearrangement of the PDZ2 domain

A systematic study of the conformational changes between the apo and holo PDZ2 scaffolds allows to gain insights into the binding mechanism. Conformational changes were first quantified in terms of the average distance between the apo and holo PDZ2 structures. To that purpose, both two-state structures were aligned with each other in UCSF Chimera (Pettersen *et al*, 2004), then the C_α_ atom coordinates were extracted and averaged over all apo and holo conformers to obtain mean apo and mean holo structures. Next, the distance between the C_α_ atom coordinates of the averaged apo and holo PDZ2 conformations was calculated for each residue. The residues for which the C_α_ deviate more than 1.5 Å from each other were highlighted and mapped on the apo PDZ2 structure as shown in Figure 3. The loop comprising residues Gly24-Gly34 and the N-terminal flexible segment were excluded from the analysis due to their elevated flexibility. Largest ligand-induced backbone conformational changes are concentrated to three sites. The first site (S1) includes α-helix 1 and part of the β-strand 2 facing it. The second site (S2) includes the middle part of the β-sheet. The third site (S3) includes parts of the β-strand 5 and α-helix 2 in proximity of the flexible loop as summarized in Figure 3. The PDZ2 allosteric network spans from the RA-GEF2 binding site including residues Val75 and Val22 to the α-helix 1 through the S1 site, to Lys54 through the S2 site, and to Val58 through the S3 site. Ligand binding yields a shift of the α-helix 1 and a part of the β-strand 2 facing it to allocate the ligand with a shift of the middle part of the β-sheet away from the binding site quantifiable also by the distance between the C_α_ atoms of residues Val22 and Val75, which is 6.5 ± 0.3 Å in the apo state 2 and 6.8 ± 0.4 Å in the apo state 1, but 7.1 ± 0.5 Å for both states in the PDZ2 complex. The aforementioned structural rearrangements are allosterically coupled to the ligand binding site, as backbone rearrangements between apo and complex PDZ2 domain are ligand induced. Those findings correlate with one of the major findings of Ranganathan et. al. who showed a statistical coupling between His71 and distal residues Ala46 and Ile52 that are part of the α-helix 1 and to the findings of Lee et. al. that indicated a coupling between residue Ile20 of the binding site and residues Ala39 and Val40 of the β-strand β3 (Fuentes *et al.*, 2004; Lockless & Ranganathan, 1999). Overall, a structural rather extensive and sophisticated correlation network of residues at atomic resolution in the coordinate space is identified that “senses” the ligand binding in line with previous indications (Fuentes *et al.*, 2004; Lockless & Ranganathan, 1999). The mechanism is of induced fit-type as already previously reported by stopped-flow measurements (Chi *et al*, 2009).

**Figure 3:**
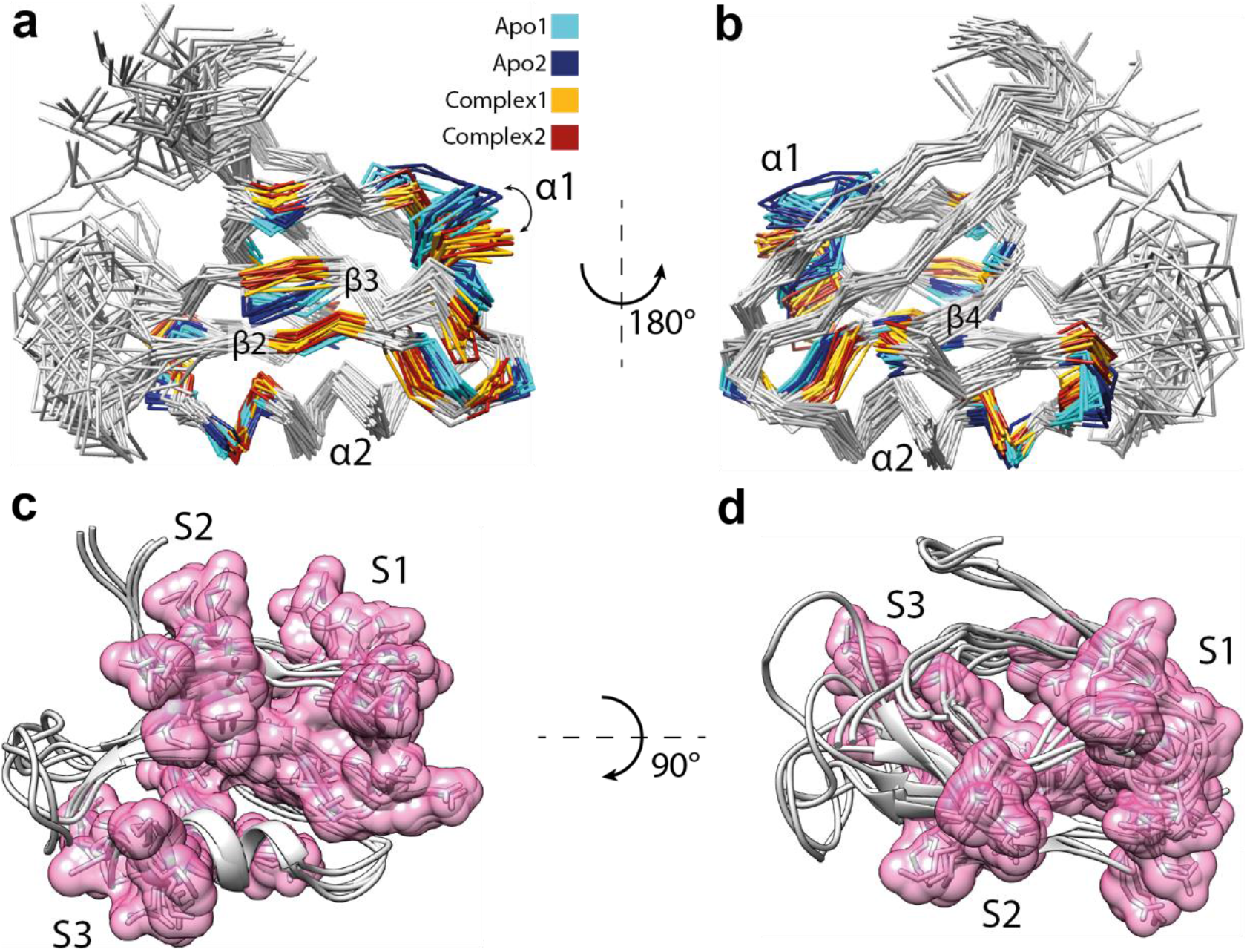
Ligand binding-induced structural changes of the PDZ2 domain. The top panel shows two views of the aligned apo and holo PDZ2 structural ensembles. All residues with distances between apo and holo averaged C_α_ positions of more than 1.5 Å except flexible regions are highlighted on the 3D protein structure with the color coding according to Figure 2 with cyan and blue representing the apo form and yellow and red the holo form, respectively (a, b). Ligand-induced allosteric movement of the α-helix 1 is indicated by the arrow. Significant rearrangements in the α-helix 1 and part of the β-strand 2 facing it, the middle part of the β-sheet and parts of the β-strand 5 and α-helix 2 in proximity of the flexible loop validate previously reported allosteric interactions in the PDZ2 domain (Fuentes *et al.*, 2004; Lockless & Ranganathan, 1999). The bottom panel shows three sites S1-S3 of the aforementioned allosteric network (c, d). The elucidated allosteric network spans from the binding site to the α-helix 1 through S1, to the Lys54 through S2, or to the Val58 through S3.

In context of multidomain proteins the induced-fit allosteric mechanism of the PDZ2 domain may connect the PDZ2 binding site with an interdomain interface. Despite the scarce information about the exact location of the potential interdomain interface for the PDZ2 domain from hPTP1E it is known that Cdc42 directly interacts with α-helix 1 from the Par-6 PDZ domain (Peterson *et al*, 2004) and a domain-domain interaction between the PDZ1 and PDZ2 occurs through a site located at the α-helix 1 / β-strand 1 region (van den Berk *et al*, 2007). This indicates a potential involvement of the α-helix 1 in the PDZ interdomain interface, that also experiences the largest ligand-induced rearrangement according to our eNOE studies upon ligand binding indicating a biological relevance of the induced-fit allostery mechanism discussed.

### Evidence for the conformational preselection in PDZ2 in terms of ligand binding

A further detailed investigation of both the apo and holo two-state structures with focus on the sidechains of residues Ala69, Thr70, His71, and Lys38 close to the binding site suggests in part a conformational selection mechanism as both holo states are overlapping with apo state 1 (Figure 4, cyan). Apo state 2 features the sidechain of residues Ala69 and His71 pointing towards the flexible loop that is pushed further away from the binding site and the sidechain of residue Lys38 pointing directly to the binding site as shown in Figure 4a (blue). The role of Lys38 was further investigated by visualization of the PDZ2 molecular surface of the two representative states from the apo and holo structures. Detailed analysis of the PDZ2 binding site conformation shows that the binding groove in the apo state 2 is obstructed by the sidechains of residues Lys38 and Lys72 as shown in Figure 5. This finding suggests state 2 has to be a “closed”, ligand-binding obstructing PDZ2 conformation while state 1 is the open, ligand-welcoming state that superimposes with the holo states. Conclusively, this indicates the presence of a conformational selection model for ligand binding.

**Figure 4:**
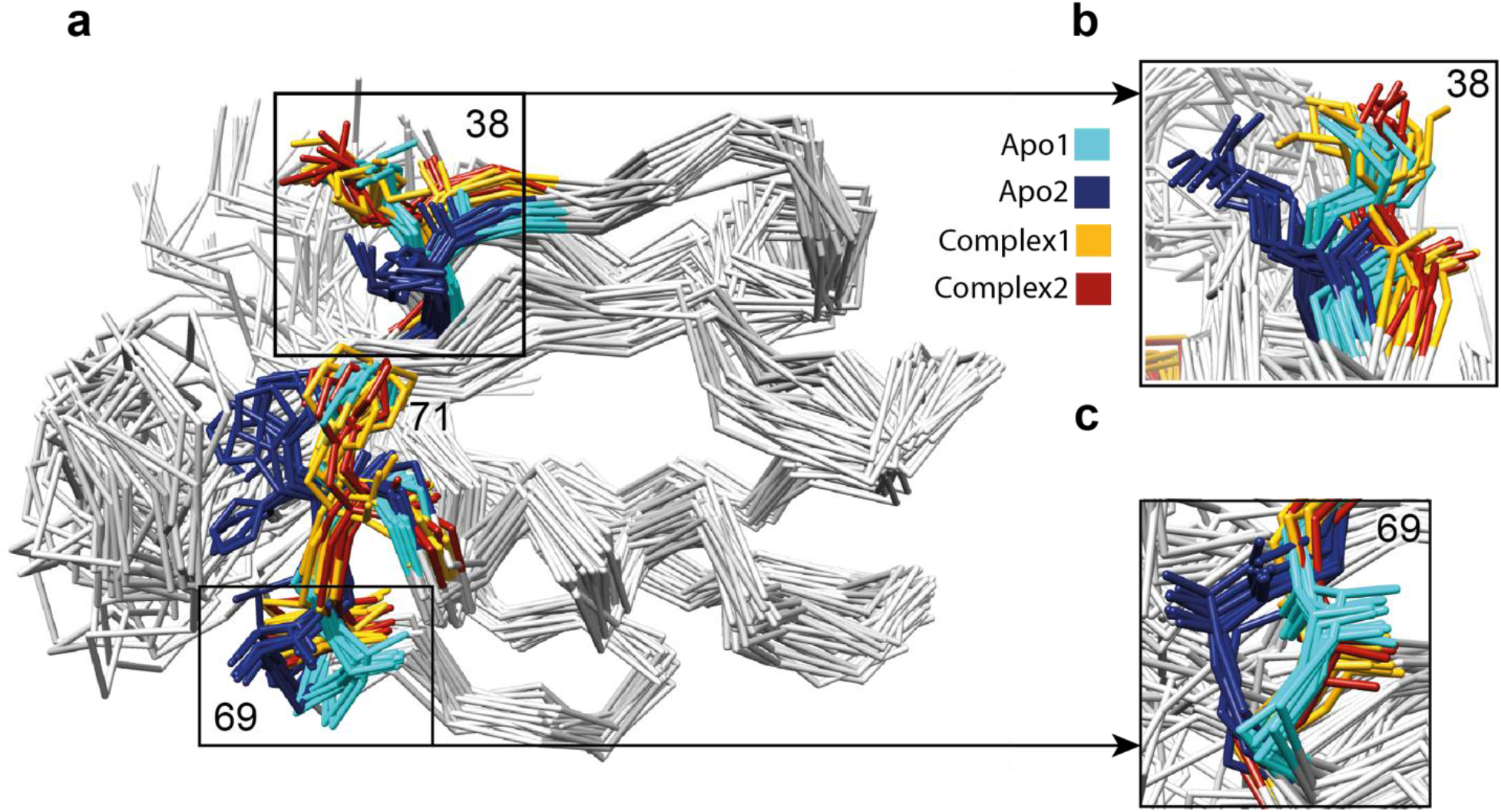
Conformational selection-based ligand binding indicated by a structure comparison of the two-state apo and holo structures. Sidechains of residues Ala69, Thr70, His71 (b) and Lys38 of the apo and holo structures color coded as in Figures 2 and 3 (a) with inserts (b and c) show a superposition of apostate 1 (cyan) with both holo states (yellow and red) suggesting a conformational selection mechanism on PDZ2 for ligand binding.

**Figure 5:**
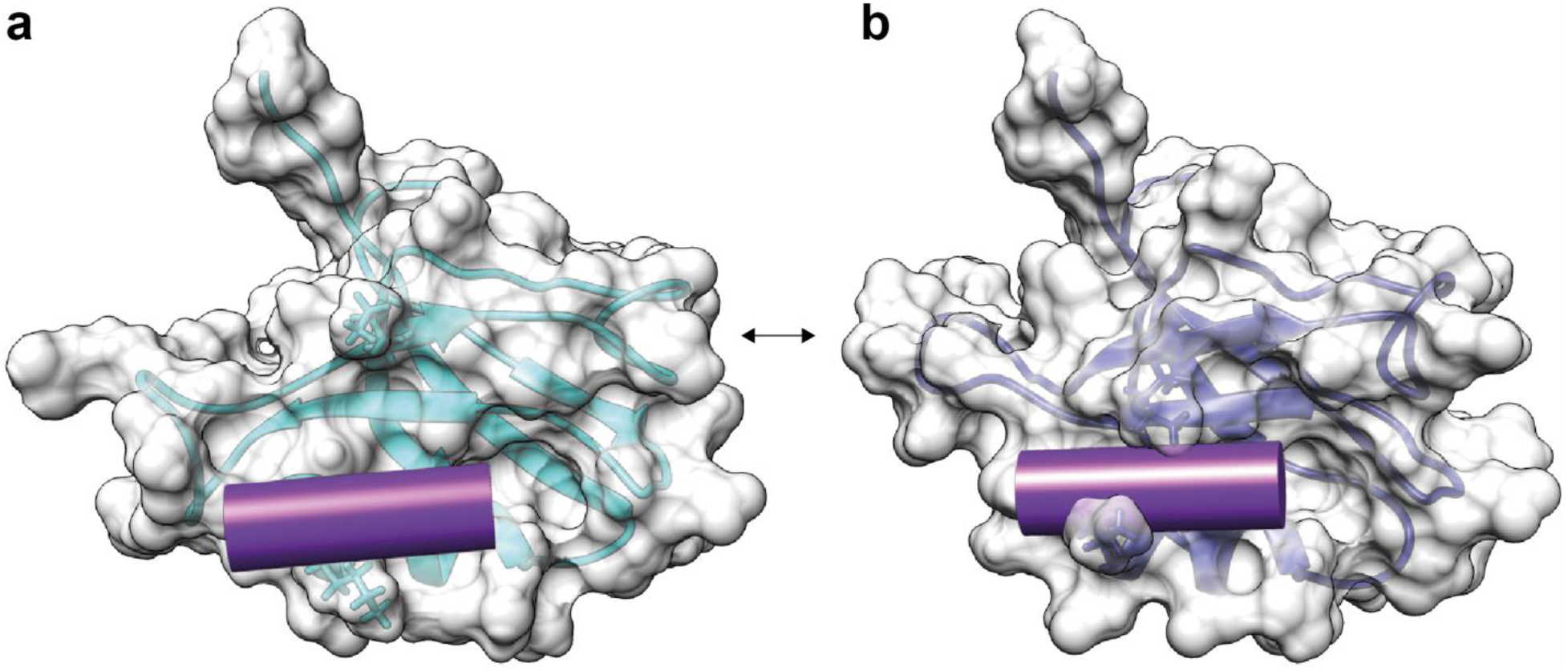
The apo form comprises a closed ligand-obstructing (a) and an open ligand-welcoming state (b). The surface views of the two states of the PDZ2 of the apo form with state 2 (a) and state 1 (b) are shown with a ribbon representation and the important side chains of Lys38 and Lys72 shining through. The position of the RA-GEF2 peptide is visualized with a violet cylinder. The access of the binding groove is obstructed in PDZ2 apo state 2 by sidechains of residues Lys38 and Lys72 (b), which hints at state preselection upon ligand binding.

### An extensive correlation network within the apo PDZ2 domain steered by the dynamic loop

The above identification of an open, ligand-welcoming and a closed state of the apo PDZ2 is now analyzed within the entire protein domain by objective extraction of correlation with the PDBcor in standard settings (Ashkinadze *et al.*). The distance correlation matrix heatmap shown in the Figure 2c indicates that apo PDZ2 is a strongly correlated protein with correlations spanning across the protein fold with exception of β-strand 1 and α-helix 1. The strongest correlations of the apo PDZ2 structural ensemble are concentrated to the RA-GEF2 binding site, β-strand 3 including residues Lys38, Lys72 and other residues involved in the conformational preselection including Ala69, Thr70 and His71 as shown in Figure 6, but ~60% of the entire protein domain is involved.

**Figure 6:**
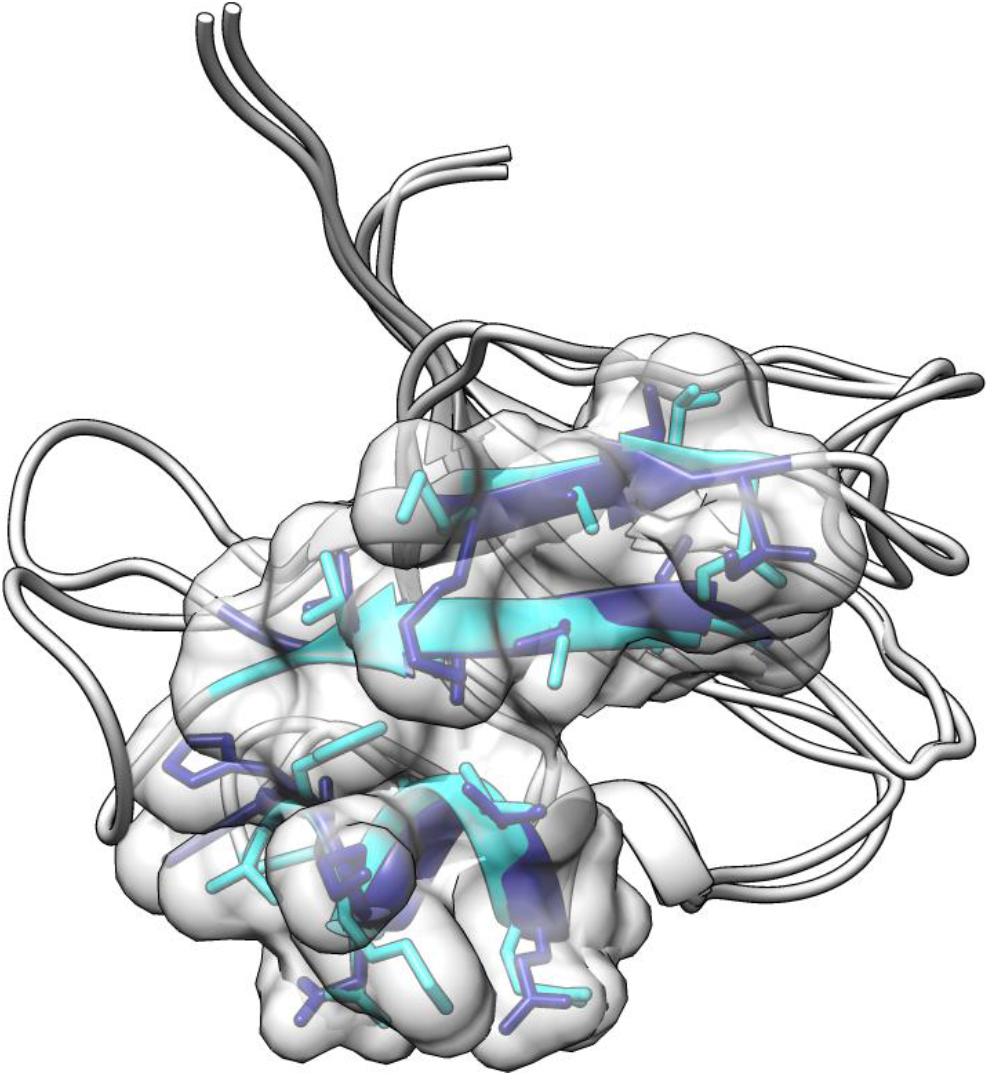
Core residues that are responsible for the two-state correlations of the apo state are localized around the ligand-binding site and the hydrophobic core of the protein. The 15 highest correlated residues as determined by the correlation matrix shown in Figure 2c are highlighted as a single volume entity on the two-state apo PDZ2 structure including state-dependent ribbon coloring and side chain representation. As it is visible from the protein 3D structure, the highest correlations are concentrated to the protein binding site and all sites involved in the conformation preselection mechanism shown Figures 4 and 5.

The analysis indicates that the presence of the two states originates from the loop comprising residues Gly24-Gly34. Due to its partial flexibility enabled by the two double glycine hinge motives Gly23-Gly24 and Gly33-Gly34 and its location at the edge of the protein structure, the loop harbors enhanced thermal intrinsic local dynamics. As a consequence, in apo state 1 the loop sterically pushes the side chains of Thr70 and His71 away enabling a shift of helix 2 closer to the loop. In addition, in state 1 a steric push of the C-terminal end of the loop along with Ile35 induces a shift of the β-sheet via Val58/Leu59.

Within this context it is interesting to note that, as in the present case of the PDZ2 domain, in the proline cis-trans isomerase human cyclophilin A a loop with two double-glycine motifs serving as hinges is key for the two-state structure of the protein comprising more than 2/3 of the protein (Chi *et al*, 2015b). While a generalization cannot be made from two cases only, the proposed mechanism of allostery that is based on a dynamic loop feed by the thermic energy of the environment, which sterically perturbs the folded part of the domain, appears to be plausible. Because of its simplicity it may be well presented in the protein world, which however remains to be demonstrated.

### On the multi-level allosteric mechanism of the PDZ2 domain

The multi-state structures of the apo and holo form of the PDZ2 domain indicate at least two levels of protein allostery. The structural correlation of the apo form comprising an extensive structural correlation network between an open ligand-welcoming and a closed ligand-obstructive state comprising ~60% of the entire domain are in line with the presence of a conformational selection allostery mechanism of ligand binding (Figure 6). In particular, the ligand-binding site with α-helix 2 and β-strand 2, and β-strand 3 are involved. Then, the ligand binding to the open state induces an extensive conformational change covering ~25% of the protein including again the binding site by definition as well as most prominently α-helix 1 and β-strand 4. Hence, with the induced fit triggers allosteric changes across the PDZ2 fold. Interestingly the two allosteric networks only partially overlap. While both networks share the binding site the apo form comprises also an allosteric network with the entire β-sheet, but α-helix 1 and β-strand 4 form another allosteric network in the holo form. Moreover, while both allosteric networks comprise the binding site they are structurally distinct also within the binding site. Finally, it is worth mentioning that the holo form comprises mainly one global structure with local plurality in the side chain configurations (i.e. distinct rotamers, Figure S5), which means that ligand-binding blocks long-range structural correlations and plasticity (highlighted in Figure 7 by the same circle shape for the yellow and red states).

**Figure 7:**
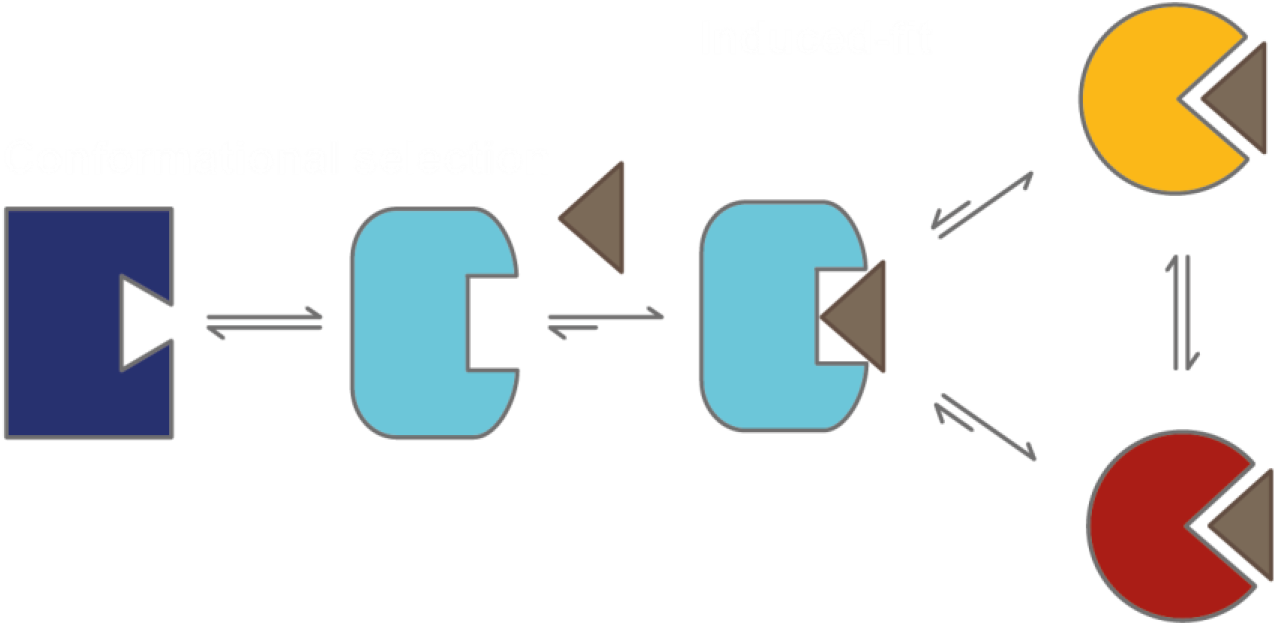
Cartoon representation of the multi-level allosteric mechanism of the PDZ2 domain. The mechanism can be summarized as a conformational selection by the ligand of the open state from an equilibrium of closed and open states of apo PDZ2, followed by an induced fit mechanism to that selected state that propagates allosteric rearrangements throughout the protein fold into the yellow and red states. The yellow and red states are distinct from each other mainly by local side chain rotamers and thus have the same shape in the cartoon. The color code of the PDZ2 domain of the Figure 2 is used and the ligand is indicated by a grey triangle.

### The various allosteric pathways within the PDZ2 domain

In a comparison between the various proposed allosteric pathways of the PDZ2 domain under study here, the methods used need to be investigated briefly. The multi-state structure determination presented shows a direct measure of a structural correlation network between states that interchange on the time scale of larger than 10 ns to micro or low milliseconds, while dynamics analysis using relaxation measurements yields a local rate between states that if similar rates are observed for parts of the protein are indicative of a correlated structural network. Dynamics can be in principle be measured from ps to milliseconds. MD simulations are usually of short time scale (i.e. in the ns range and for the example of interest PDZ2 domain 16 replicas of a 2 ns MD trajectory calculation were performed (Dhulesia *et al.*, 2008)). The evolutionary approach differs from the others as it does not only evaluate a single hPTP1E PDZ2 domain, but a whole family of the PDZ domains. This makes this approach sensitive to a conserved allosteric interaction among all PDZ domains. Indeed, the allosteric pathway deduced from evolutionary data of the PDZ domain (Lockless & Ranganathan, 1999) resembles our eNOE-based induced-fit allosteric network since it connects the binding site with α-helix 1 as it can be seen from the comparison between allosteric networks shown in Figure 8. In contrast, the allosteric pathways deduced from MD data (Gerek & Ozkan, 2011) and methyl relaxation data (Fuentes *et al.*, 2004) resemble rather our eNOE-based conformational selection allosteric network covering the PDZ binding site and β-strands 2 and 3 (Figure 8).

**Figure 8:**
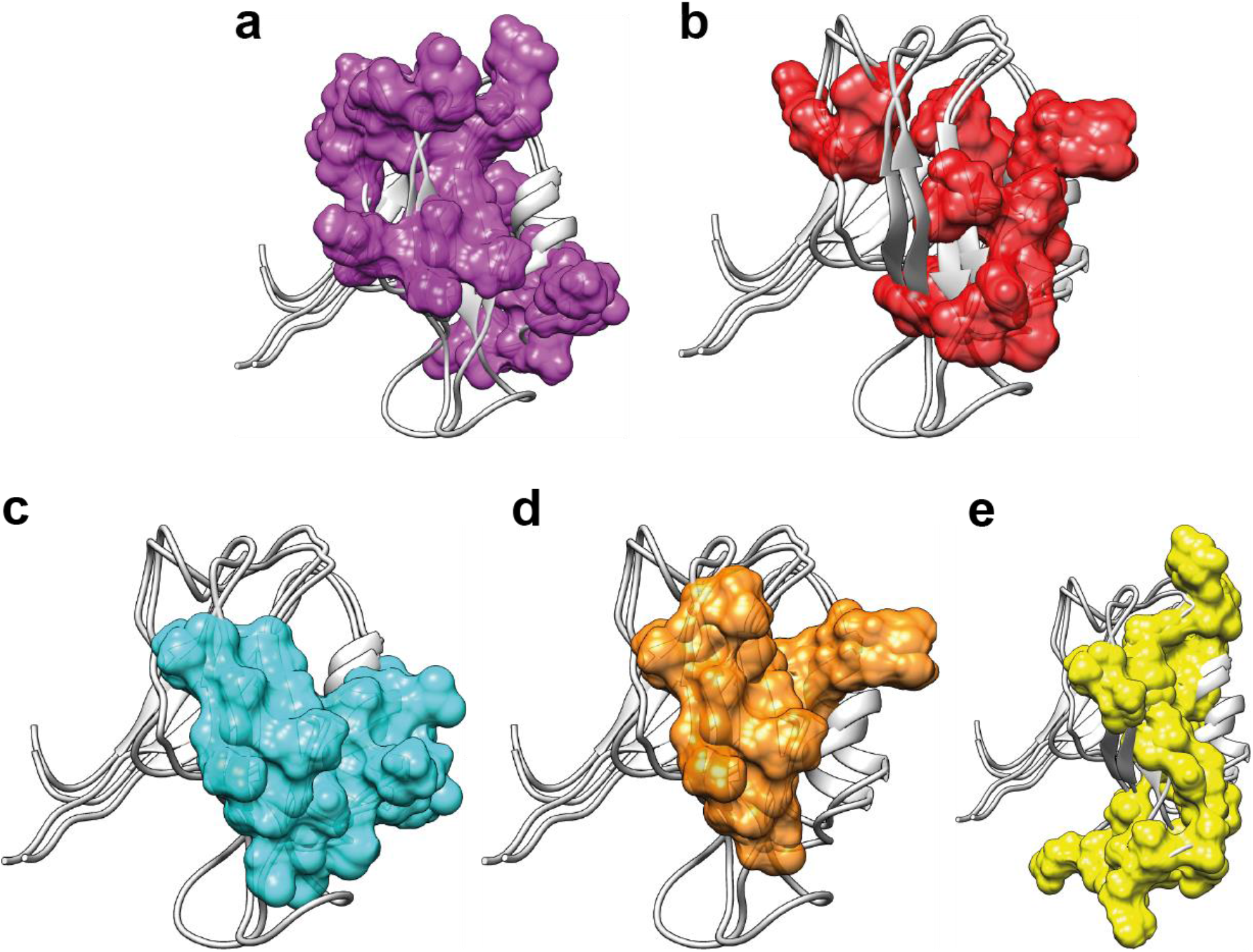
Proposed allosteric pathways within PDZ2 domain determined or predicted with various methods including eNOE-based induced-fit allosteric network (a), allosteric network based on evolutionary coupling in the family of the PDZ domains (b; (Lockless & Ranganathan, 1999)), eNOE-based conformation-selection allosteric network of the free PDZ2 domain (c), allosteric network based on MD simulation (d; (Gerek & Ozkan, 2011)), and allosteric network based on methyl relaxation data (e; (Fuentes *et al.*, 2004)).

## CONCLUSION

The presented work on the well-studied PDZ2 domain showcases the power of NMR with the high accuracy afforded by eNOEs that gives the possibility to solve multiple protein states at atomic resolution under physiological conditions for the studies of protein allostery. It elucidated a two-level allosteric network at atomic resolution and pinpoints to the existence of rather extensive and sophisticated structural correlation networks of residues that in principle could be used for ligand binding regulation or signaling. In the context of the system of interest, the PDZ2 allosteric binding mechanism was found to be combined from the broadly accepted induced-fit and conformational selection mechanisms. In more general terms, the presented work validates also in part previously reported allosteric indicators using a genetic algorithm (Lockless & Ranganathan, 1999), experimental data (Fuentes *et al.*, 2004) and MD simulation (Gerek & Ozkan, 2011). Furthermore, such properties are to be expected in almost any biomolecular system awaiting to get explored.

## DEPOSITED PROTEIN STRUCTURES

Following PDB codes were deposed at Protein Data Bank: XXXX (apo PDZ2) and XXXX (PDZ2 bound to RA-GEF2 peptide)

## ACKNOWLEDGMENT

We would like to thank the Swiss National Science Foundation (SNF), the National Science Foundation (NSF grant 1917254 for Infrastructure Innovation for Biological Research to B.V.) and the National Institutes of Health (NIH grant R01GM130694-01 to B.V.) for financial support.

## REFERENCES

Ashkinadze D, Klukowski P, Kadavath H, Güntert P, Riek R PDBcor: An Automated Correlation Extraction Calculator for Multi-State Protein Structures. Available at SSRN 3904349

Bai Y, Englander SW (1996) Future directions in folding: The multi‐state nature of protein structure. Proteins: Structure, Function, and Bioinformatics 24: 145–151

Born A, Soetbeer J, Breitgoff F, Henen MA, Sgourakis N, Polyhach Y, Nichols PJ, Strotz D, Jeschke G, Vögeli B (2021) Reconstruction of Coupled Intra-and Interdomain Protein Motion from Nuclear and Electron Magnetic Resonance. Journal of the American Chemical Society

Chi CN, Bach A, Engström Å, Wang H, Strømgaard K, Gianni S, Jemth P (2009) A sequential binding mechanism in a PDZ domain. Biochemistry 48: 7089–7097

Chi CN, Strotz D, Riek R, Vögeli B (2015a) Extending the eNOE data set of large proteins by evaluation of NOEs with unresolved diagonals. Journal of biomolecular NMR 62: 63–69

Chi CN, Vögeli B, Bibow S, Strotz D, Orts J, Güntert P, Riek R (2015b) A structural ensemble for the enzyme cyclophilin reveals an orchestrated mode of action at atomic resolution. Angewandte Chemie International Edition 54: 11657–11661

Clore GM, Starich MR, Bewley CA, Cai M, Kuszewski J (1999) Impact of residual dipolar couplings on the accuracy of NMR structures determined from a minimal number of NOE restraints. Journal of the American Chemical Society 121: 6513–6514

Cooper A, Dryden D (1984) Allostery without conformational change. European Biophysics Journal 11: 103–109

Dalvit C, Fogliatto G, Stewart A, Veronesi M, Stockman B (2001) WaterLOGSY as a method for primary NMR screening: practical aspects and range of applicability. Journal of biomolecular NMR 21: 349–359

Dhulesia A, Gsponer J, Vendruscolo M (2008) Mapping of two networks of residues that exhibit structural and dynamical changes upon binding in a PDZ domain protein. Journal of the American Chemical Society 130: 8931–8939

Doyle DA, Lee A, Lewis J, Kim E, Sheng M, MacKinnon R (1996) Crystal structures of a complexed and peptide-free membrane protein–binding domain: molecular basis of peptide recognition by PDZ. Cell 85: 1067–1076

Emmanouilidis L, Schütz U, Tripsianes K, Madl T, Radke J, Rucktäschel R, Wilmanns M, Schliebs W, Erdmann R, Sattler M (2017) Allosteric modulation of peroxisomal membrane protein recognition by farnesylation of the peroxisomal import receptor PEX19. Nature communications 8: 1–13

Fenwick RB, Esteban-Martín S, Richter B, Lee D, Walter KF, Milovanovic D, Becker S, Lakomek NA, Griesinger C, Salvatella X (2011a) Weak long-range correlated motions in a surface patch of ubiquitin involved in molecular recognition. Journal of the American Chemical Society 133: 10336–10339

Fenwick RB, Esteban-Martín S, Salvatella X (2011b) Understanding biomolecular motion, recognition, and allostery by use of conformational ensembles. European Biophysics Journal 40: 1339–1355

Fenwick RB, Orellana L, Esteban-Martín S, Orozco M, Salvatella X (2014) Correlated motions are a fundamental property of β-sheets. Nature communications 5: 1–9

Fenwick RB, Schwieters CD, Vogeli B (2016) Direct investigation of slow correlated dynamics in proteins via dipolar interactions. Journal of the American Chemical Society 138: 8412–8421

Frauenfelder H, McMahon BH, Austin RH, Chu K, Groves JT (2001) The role of structure, energy landscape, dynamics, and allostery in the enzymatic function of myoglobin. Proceedings of the National Academy of Sciences 98: 2370–2374

Fuentes EJ, Der CJ, Lee AL (2004) Ligand-dependent dynamics and intramolecular signaling in a PDZ domain. Journal of molecular biology 335: 1105–1115

Gerek ZN, Ozkan SB (2011) Change in allosteric network affects binding affinities of PDZ domains: analysis through perturbation response scanning. PLoS computational biology 7: e1002154

Green SM, Shortle D (1993) Patterns of nonadditivity between pairs of stability mutations in staphylococcal nuclease. Biochemistry 32: 10131–10139

Güntert P (2004) Automated NMR structure calculation with CYANA. In: Protein NMR Techniques, pp. 353–378. Springer:

Güntert P, Buchner L (2015) Combined automated NOE assignment and structure calculation with CYANA. Journal of biomolecular NMR 62: 453–471

Güntert P, Mumenthaler C, Wüthrich K (1997) Torsion angle dynamics for NMR structure calculation with the new program DYANA. Journal of molecular biology 273: 283–298

Harris BZ, Lim WA (2001) Mechanism and role of PDZ domains in signaling complex assembly. Journal of cell science 114: 3219–3231

Hung AY, Sheng M (2002) PDZ domains: structural modules for protein complex assembly. Journal of Biological Chemistry 277: 5699–5702

Ishima R, Torchia DA (2000) Protein dynamics from NMR. Nature structural biology 7: 740–743

Iwahara J, Schwieters CD, Clore GM (2004) Ensemble approach for NMR structure refinement against 1H paramagnetic relaxation enhancement data arising from a flexible paramagnetic group attached to a macromolecule. Journal of the American Chemical Society 126: 5879–5896

Karplus M, Weaver DL (1976) Protein-folding dynamics. Nature 260: 404–406

Keepers JW, James TL (1984) A theoretical study of distance determinations from NMR. Two-dimensional nuclear Overhauser effect spectra. Journal of Magnetic Resonance (1969) 57: 404–426

Kozlov G, Banville D, Gehring K, Ekiel I (2002) Solution structure of the PDZ2 domain from cytosolic human phosphatase hPTP1E complexed with a peptide reveals contribution of the β2– β3 loop to PDZ domain–ligand interactions. Journal of molecular biology 320: 813–820

Kozlov G, Gehring K, Ekiel I (2000) Solution structure of the PDZ2 domain from human phosphatase hPTP1E and its interactions with C-terminal peptides from the Fas receptor. Biochemistry 39: 2572–2580

Kumar A, 1985. Two-dimensional nuclear Overhauser effect in biomolecules, Proceedings of the Indian Academy of Sciences-Chemical Sciences. Springer, pp. 1–8.

Lockless SW, Ranganathan R (1999) Evolutionarily conserved pathways of energetic connectivity in protein families. Science 286: 295–299

Monnot C, Bihoreau C, Conchon S, Curnow KM, Corvol P, Clauser E (1996) Polar residues in the transmembrane domains of the type 1 angiotensin II receptor are required for binding and coupling: reconstitution of the binding site by co-expression of two deficient mutants. Journal of Biological Chemistry 271: 1507–1513

Nichols PJ, Born A, Henen MA, Strotz D, Orts J, Olsson S, Güntert P, Chi CN, Vögeli B (2017) The exact nuclear overhauser enhancement: recent advances. Molecules 22: 1176

Nichols PJ, Henen MA, Born A, Strotz D, Güntert P, Vögeli B (2018) High-resolution small RNA structures from exact nuclear Overhauser enhancement measurements without additional restraints. Communications biology 1: 1–11

Orts J, Vögeli B, Riek R (2012) Relaxation matrix analysis of spin diffusion for the NMR structure calculation with eNOEs. Journal of chemical theory and computation 8: 3483–3492

Palmer A, Zimmer M, Erdmann KS, Eulenburg V, Porthin A, Heumann R, Deutsch U, Klein R (2002) EphrinB phosphorylation and reverse signaling: regulation by Src kinases and PTP-BL phosphatase. Molecular cell 9: 725–737

Peng JW (2015) Investigating dynamic interdomain allostery in Pin1. Biophysical reviews 7: 239–249

Peterson FC, Penkert RR, Volkman BF, Prehoda KE (2004) Cdc42 regulates the Par-6 PDZ domain through an allosteric CRIB-PDZ transition. Molecular cell 13: 665–676

Petit CM, Zhang J, Sapienza PJ, Fuentes EJ, Lee AL (2009) Hidden dynamic allostery in a PDZ domain. Proceedings of the National Academy of Sciences 106: 18249–18254

Pettersen EF, Goddard TD, Huang CC, Couch GS, Greenblatt DM, Meng EC, Ferrin TE (2004) UCSF Chimera—a visualization system for exploratory research and analysis. Journal of computational chemistry 25: 1605–1612

Rod TH, Radkiewicz JL, Brooks CL (2003) Correlated motion and the effect of distal mutations in dihydrofolate reductase. Proceedings of the National Academy of Sciences 100: 6980–6985

Sato T, Irie S, Kitada S, Reed JC (1995) FAP-1: a protein tyrosine phosphatase that associates with Fas. Science 268: 411–415

Selvaratnam R, Chowdhury S, VanSchouwen B, Melacini G (2011) Mapping allostery through the covariance analysis of NMR chemical shifts. Proceedings of the National Academy of Sciences 108: 6133–6138

Strotz D, Orts J, Chi CN, Riek R, Vögeli B (2017) ENORA2 exact NOE analysis program. Journal of chemical theory and computation 13: 4336–4346

Strotz D, Orts J, Kadavath H, Friedmann M, Ghosh D, Olsson S, Chi CN, Pokharna A, Güntert P, Vögeli B (2020) Protein allostery at atomic resolution. Angewandte Chemie International Edition 59: 22132–22139

Swain JF, Gierasch LM (2006) The changing landscape of protein allostery. Current opinion in structural biology 16: 102–108

Tonikian R, Zhang Y, Sazinsky SL, Currell B, Yeh J-H, Reva B, Held HA, Appleton BA, Evangelista M, Wu Y (2008) A specificity map for the PDZ domain family. PLoS biology 6: e239

van den Berk LC, Landi E, Walma T, Vuister GW, Dente L, Hendriks WJ (2007) An Allosteric Intramolecular PDZ− PDZ Interaction Modulates PTP-BL PDZ2 Binding Specificity. Biochemistry 46: 13629–13637

Vögeli B (2014) The nuclear Overhauser effect from a quantitative perspective. Progress in nuclear magnetic resonance spectroscopy 78: 1–46

Vögeli B, Kazemi S, Güntert P, Riek R (2012) Spatial elucidation of motion in proteins by ensemble-based structure calculation using exact NOEs. Nature structural & molecular biology 19: 1053–1057

Vögeli B, Olsson S, Güntert P, Riek R (2016) The exact NOE as an alternative in ensemble structure determination. Biophysical journal 110: 113–126

Vögeli B, Segawa TF, Leitz D, Sobol A, Choutko A, Trzesniak D, van Gunsteren W, Riek R (2009) Exact distances and internal dynamics of perdeuterated ubiquitin from NOE buildups. Journal of the American Chemical Society 131: 17215–17225

Vögeli B, Vugmeyster L (2019) Distance‐independent cross‐correlated relaxation and isotropic chemical shift modulation in protein dynamics studies. ChemPhysChem 20: 178–196

Volkman BF, Lipson D, Wemmer DE, Kern D (2001) Two-state allosteric behavior in a single-domain signaling protein. Science 291: 2429–2433

Walma T, Spronk CA, Tessari M, Aelen J, Schepens J, Hendriks W, Vuister GW (2002) Structure, dynamics and binding characteristics of the second PDZ domain of PTP-BL. Journal of molecular biology 316: 1101–1110

Wüthrich K (1990) Protein structure determination in solution by NMR spectroscopy. Journal of Biological Chemistry 265: 22059–22062

